# Solution AFM imaging and coarse-grained molecular modeling of yeast condensin structural variation coupled to the ATP hydrolysis cycle

**DOI:** 10.1101/2024.12.16.628603

**Authors:** Hiroki Koide, Noriyuki Kodera, Shoji Takada, Tsuyoshi Terakawa

## Abstract

Condensin is a protein complex that regulates chromatin structural changes during mitosis. It varies the molecular conformation through the ATP hydrolysis cycle and extrudes DNA loops into its ring-like structure as a molecular motor. Condensin contains Smc2 and Smc4, in which a coiled-coil arm tethers the hinge and head domains and dimerizes at the hinge. ATPs bind between the heads, induce their engagement, and are hydrolyzed to promote their disengagement. In the previous study, we performed solution atomic force microscopy (AFM) imaging of yeast condensin holo-complex in the presence of ATP and conducted flexible molecular fitting to the AFM image, obtaining the hinge structure with open conformation. However, it has yet to be clarified how the opening/closing of the hinge is coupled to the ATP hydrolysis cycle. In this study, we performed solution AFM imaging in the presence and absence of varying nucleotides, including AMP-PNP, ATPγS, and ADP. Furthermore, we conducted coarse-grained molecular dynamics simulations of a Smc2/4 heterodimer and selected the molecular structure that best represented each AFM image. Our results suggested that head engagement upon ATP binding is coupled to hinge opening. Also, the results indicated that the N-terminal region of Brn1, one of the accessory subunits, re-associates to the Smc2 head after ADP release. This study deepens our understanding of the conformational variation of yeast condensin driven by the ATP hydrolysis cycle.

## INTRODUCTION

Chromosome condensation and proper segregation are crucial processes during cell division, and dysfunction can lead to uneven distribution of genetic information and cell death. (1–6). Condensin is a protein complex that governs the structural changes of chromatin, particularly during mitosis, and plays vital roles in promoting chromatin condensation and supporting accurate chromosome segregation (7–9). Condensin also dynamically changes its molecular structure through the ATP hydrolysis cycle and acts as a molecular motor to extrude DNA loops into its circular structure to expand it, thereby achieving chromosome condensation (10–14). In recent years, molecular mechanisms other than loop extrusion, such as phase separation, have also been proposed (15–17).

Analysis of the molecular structure of condensin has revealed that each monomer of Smc2 and Smc4 has hinge and head domains, which are connected via a coiled-coil arm region (18–21). The hinge is involved in the formation of Smc2/4 dimers. Meanwhile, ATP binds to the interface between the heads of Smc2/4, and this ATP binding induces engagement of the two heads. When ATP is hydrolyzed, nucleotides are released, and the heads disengage (22, 23). Atomic force microscopy (AFM) studies have revealed that the Smc2/4 heterodimer can alternate between an O-form in which the hinge and heads are engaged and the arms are widely opened, a B-form in which the hinge and heads are close to each other, a V-form in which the heads are disengaged, and an I-form in which the coiled-coil regions are closely aligned (24, 25). However, only the I-form-like structure realized when nucleotides are not bound to the heads has been elucidated in structural characterization by cryo-electron microscopy (EM) (19).

The X-ray crystal structures of parts of condensin have also been solved. For example, closed hinge structures of eukaryotic and archaeal condensins, in which two partial coiled-coil arms protruding from the hinge are closely aligned, have been obtained (26, 27). On the other hand, an open hinge structure of bacterial condensins in which arms are widely opened has also been solved (26). In a previous study, we performed solution AFM imaging of the budding yeast condensin holo-complex in the presence of ATP and demonstrated that an open hinge structure must be assumed to fit the molecular structure to the hinge part of the observed O-form condensin (28). However, it is unclear how the transition from the closed to the open hinge structure is linked to the nucleotide-binding state of the heads and their engage-to-disengage dynamics.

In the condensin holo-complex, the N- and C-termini of the intrinsically disordered kleisin subunit Brn1 are bound to the heads of Smc2 and Smc4, respectively, and the HEAT repeat subunits Ycg1 and Ycs4 are bound to the Brn1 (18–21). Previous biochemical experiments have shown that the N-terminus of Brn1 dissociates from Smc2 upon two ATP molecules binding to the heads (22). They introduced a protease cleavage site into Brn1 and evaluated the binding state of the cleaved Brn1 before and after adding ATP. As a result, the interaction between the Smc2 head and N-terminus of Brn1 was disrupted, and the Brn1 was dissociated in the presence of ATP. This result indicated that ATP-dependent engagement of the Smc2/4 heads causes the dissociation of Brn1 and the opening of the kleisin compartment. However, the ATP hydrolysis cycle stage at which the reassociation of Brn1 and the Smc2 head happens remains unclear.

In this study, we conducted solution AFM imaging of the condensin holo-complex in the presence and absence of varying nucleotides, including AMP-PNP, ATPγS, and ADP, to understand the molecular dynamics coupled to the ATP hydrolysis cycle. Furthermore, we performed coarse-grained molecular dynamics simulations in which the potential energy that stabilizes the combination of open/closed hinge structures and engaged/disengaged head structures was imposed. Then, the structure that best represents each AFM image was selected from the obtained structural ensemble. As a result, we found that condensin adopts One-form, O-form, I-form, V-form, and Hat-form, depending on the nucleotide state. The result also indicated that nucleotide association induces the open hinge structure, and the nucleotide dissociation closes the hinge. In addition, the Hat-form, in which Brn1 and Smc2 might be dissociated, was enriched in the presence of ADP, suggesting that reassociation of Brn1 and Smc2 happens after ADP release. The results of this study will deepen our understanding of the mechanism of conformational changes in condensin driven by the ATP hydrolysis cycle.

## RESULTS

### Solution AFM imaging of condensin in the presence and absence of various nucleotides

To clarify how the molecular conformation of the budding yeast condensin holo-complex varies depending on the nucleotide state, we performed solution AFM measurements in the presence and absence of ATP, ATPγS, AMP-PNP, and ADP. ATPγS is an ATP analog that is hydrolyzed 100 times slower than ATP, and AMP-PNP is a non-hydrolyzable ATP analog. The observed representative images are shown in Figure 1A and Figure S1. In the absence of nucleotides, we observed a small globular domain (385 nm^3^), a large globular domain (2540 nm^3^), and a string-like region connecting them. These are thought to be the dimerized hinge, a complex of the engaged heads and HEAT subunits (Ycg1 and Ycs4), and coiled-coil arms, respectively. The length of the coiled-coil region is approximately 39 nm, which roughly corresponds to the length between the head and the elbow in the cryo-EM structure in the absence of nucleotides (32 nm) (19). In some images, we also observed kinking of the elbow (Figure 1B). These structures are reminiscent of the cryo-EM structure of condensin in the absence of nucleotides (19). We call this conformation the One-form hereafter. In some images, another globular moiety was observed in addition to the complex of the heads and the HEAT subunits, and when this globular moiety appeared, the volume of the heads–HEAT subunit complex seemed to become smaller. This observation is consistent with the results of a previous cryo-EM study and is thought to be Ycg1 dissociated from the heads (Figure 1A) (19).

**Figure 1.**
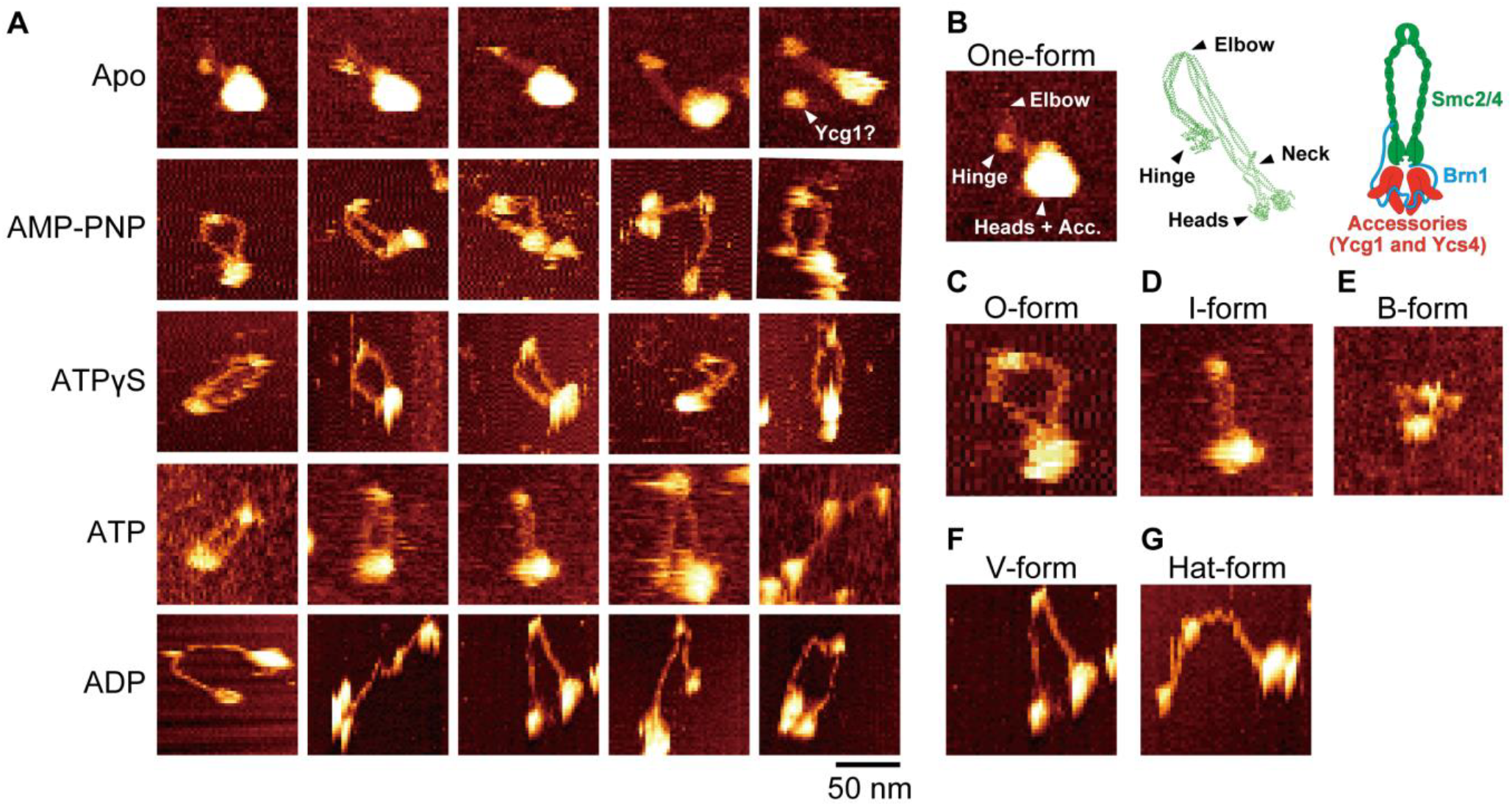
Solution AFM imaging of budding yeast condensin holo-complex. (A) Example images in the Apo (in the absence of nucleotides), AMP-PNP, ATPγS, ATP, and ADP states. (B) The representative image of the One-form, the cryo-EM structure in the Apo state (PDB ID: 6YVU) (19), and the schematic of the budding yeast condensin holo-complex. (C–G) The representative images of the (C) O-, (D) I-, (E) B-, (F) V-, and (G) Hat-forms.

On the other hand, when ATPγS or AMP-PNP was added, the two arms connecting the hinge and heads were observed to be widely open (Figure 1C). This result is consistent with previous AFM observations that showed the formation of the O-form in the presence of ATP (24, 25). In addition, in some images, the B-form was observed in which the hinge folds back to a complex of heads and HEAT subunits (Figure 1E). This result is also consistent with the previous AFM observation (25). However, while the B-form was observed in about 40–50% of the images in the previous study (25), it was observed in 4% of the images (N = 7) in the current study. This difference may be due to the difference in buffer conditions [previous study (25): 20 mM Tris-HCl pH 7.5, 50 mM NaCl, 2.5 mM MgCl_2_, 2.5 mM DTT; this study: 40 mM Tris-HCl pH7.5, 125 mM KCl, 10 mM MgCl_2_, 1 mM DTT]. Notably, under the buffer conditions in this study, the One-form was observed in almost 100% of the condensin molecules without nucleotides (N = 64). In the previous study, the I-form (Figure. 1D) without elbow kinking was observed in about 10% of the condensin molecules without nucleotides (25). Interestingly, a dramatic resolution improvement in the AFM images was observed by changing the monovalent salt from NaCl to KCl and increasing the MgCl_2_ to 10 mM, as previously reported (29).

Next, we performed solution AFM imaging in the presence of ADP to obtain information on the condensin conformation after ATP hydrolysis. As far as we know, this is the first time that AFM imaging of the budding yeast condensin holo-complex has been performed in the presence of ADP. As a result, we observed many V-form condensins with disengaged heads (Figure 1A, F, G). The One-form, which was dominant in the absence of nucleotides, was hardly observed, suggesting that the nucleotide-binding site is saturated by 4 mM ADP. Interestingly, we also found that some of the V-form structures had head-hinge-head angles of less than 90° (Figure 1F), while others were widely opened at more than 90° (Figure 1G) (we call them V-form and Hat-form, respectively, hereafter). To clarify whether these conformations represent distinct states with free energy barriers for the transition between them, we need to perform molecular structural fitting and ensemble analysis, as described below.

Next, the AFM imaging of condensin in the presence of ATP revealed the I-form, O-form, B-form, V-form, and Hat-form structures (Figure 1A). This suggests that the ATP hydrolysis cycle is linked to the transitions among these structures. Interestingly, the I-form (Figure 1D) observed in the presence of ATP differs from the One-form (Figure 1B), which is dominant in the absence of nucleotides, in that the elbow region is not bent. This result suggests almost no genuine One-form in 4 mM ATP.

### Coarse-grained molecular dynamics simulations of the Smc2/4 dimer

To generate a molecular structure for fitting to each AFM image, we next performed coarse-grained molecular dynamics simulations of the Smc2/4 dimer using the AICG2+ model (30, 31). In this model, coarse-grained beads were first placed at the Cα atom positions of each amino acid in the native all-atom structure of the protein. Then, potential energy functions that stabilize the bond lengths and non-local contacts in the native reference structure were imposed between amino acids. The potential energy functions were also imposed based on the statistics of bond angles and dihedral angles in the loop regions of proteins registered in the Protein Data Bank (https://www.rcsb.org/) (32). Also, an excluded volume energy function was imposed on amino acid pairs that were not in contact with each other in the native structure, and protein structures were time-propagated by the Langevin equation (33). The parameters of the AICG2+ model were determined to reproduce the fluctuations around the representative native protein structures (30). The elbows of the coiled-coil arms were treated specially. First, the residues in the arms of the cryo-EM structure (PDB ID: 6YVU) (19) were assigned to form either an α-helix, a β-sheet, or a loop using DSSP software (34). As a result, several residues in Smc2 (S383, T384, L385, A395, D396, G397, G398, Y785, D786, S787, S791, K792) and Smc4 (S539, L540, K541, D542, K543, M967, K968, I969, I972) were assigned to form loops (red regions in Figures 2A). Therefore, we did not impose a potential energy function to stabilize non-local contacts involving these residues. This treatment allowed us to sample structures where the elbow was not bent, as observed in the solution AFM imaging (24, 25, 28).

**Figure 2.**
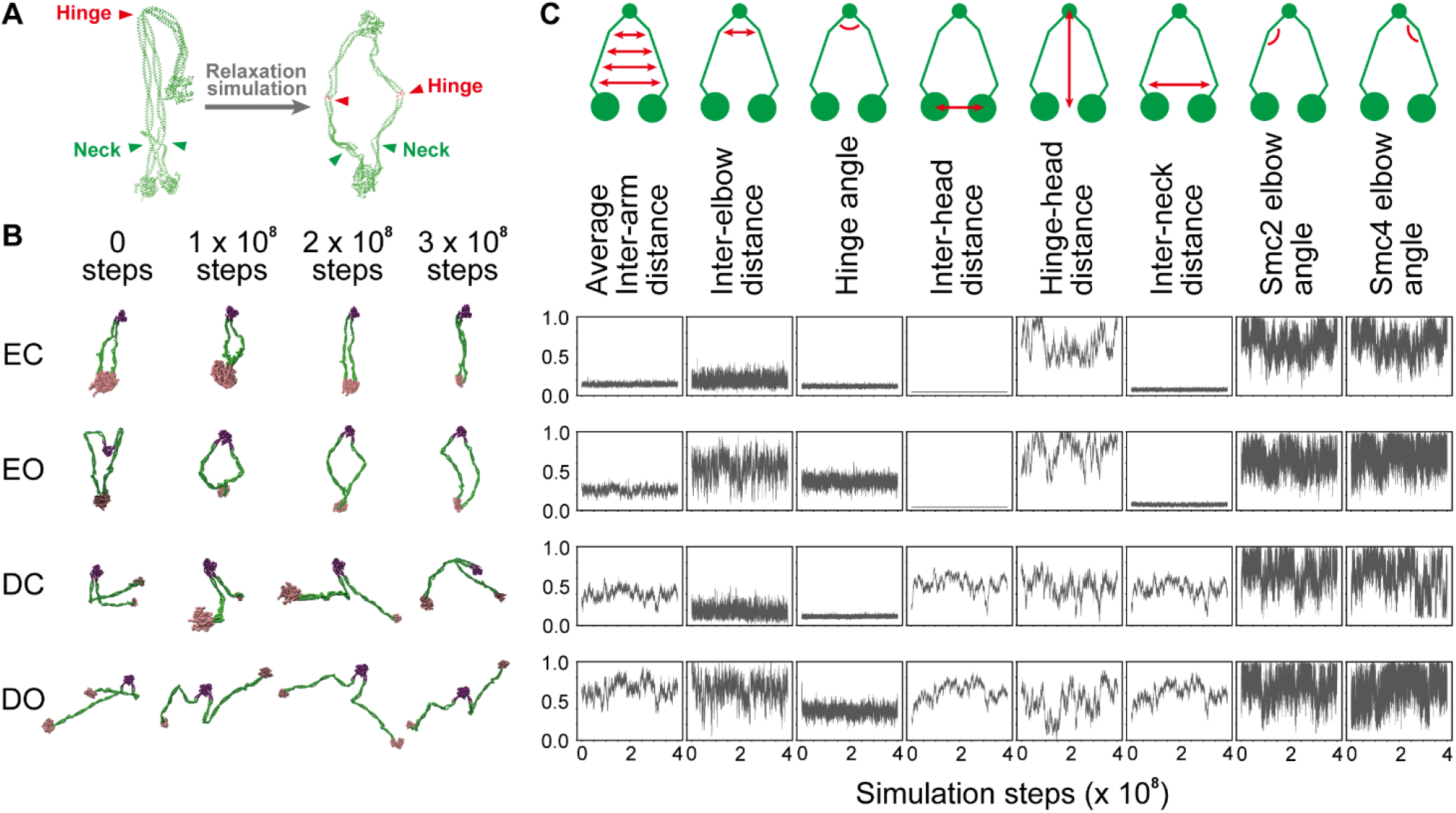
Coarse-grained molecular dynamics simulations of the SMC2/4 dimer. (A) The schematic of the relaxation simulations. (B) The snapshots from the simulations in each condition. The hinge, arms, and heads are colored purple, green, and pink, respectively. (C) The time trajectories of the 8 features defined in Methods from the simulations in each condition.

Previous studies have shown that the Smc2/4 heads transition between engaged and disengaged states (22, 23). Also, it has been suggested that the hinge transition between an open and a closed structure (28). In this study, we combined these findings and performed simulations in the four conditions: engaged heads + open hinge (EO), engaged heads + closed hinge (EC), disengaged heads + open hinge (DO), and disengaged heads + closed hinge (DC) (Figure 2B). In the simulations to stabilize the closed hinge, we used the hinge structure in One-form (PDB ID: 6YVU) (19) as the native reference structure and imposed a potential energy function stabilizing non-local contacts in the reference structure. On the other hand, in the simulations to stabilize the open hinge, we used the all-atom structure, which MODELLER (35) generated using the coarse-grained open hinge structure previously obtained by flexible fitting to an AFM image (28), as a native reference structure. In these simulations, the native contact pairs formed between the coiled-coil arms in the initial One-form structure (PDB ID: 6YVU) (19) were removed from the non-local contact list. The heads in 6YVU (19) were used as the reference structure for the simulation of the disengaged state. On the other hand, the heads in 6YVD (19) were used as the reference structure for the simulation of the engaged state. In each condition, a 1 × 10^7^ step relaxation was performed at 300 K (Figure 2A), followed by 25 production runs of 4 × 10^8^ steps at 300 K (Figure 2B). The structural coordinates were saved every 10^4^ steps, and the first 15,000 structures were excluded from the ensemble to minimize the influence of the initial structure.

As a result, we obtained a structural ensemble containing various structures in the conditions EC, EO, DC, and DO (Figure 2C), each containing 625,000 structures. The hinge-head distance, Smc2 elbow angle, and Smc4 elbow angle (Methods for definitions) fluctuated wildly in all the conditions. This result suggested that the elbow can bend flexibly regardless of the engagement of the head and the opening/closing of the hinge. In the EC condition, the inter-arm average distance (average value: 56.1± 6.7Å), inter-elbow distance (56.1±15.8Å), inter-neck distance (56.6±7.7Å), inter-head distance (41.7±0.5Å), and hinge angle (20.6±2.0°) (Methods for definitions) all fluctuated around small values. In the EO condition, the inter-elbow distance (160.5±40.1Å) and hinge angle (66.0±9.1°) fluctuated around relatively large values. In the DC condition, the average inter-arm distance (151.9±27.0Å), inter-neck distance (356.5±70.4Å), and inter-head distance (492±90.1Å) fluctuated around relatively large values. All values fluctuated around relatively large values under the DO condition. These results suggested that non-overlapping structural ensembles were sampled in the simulations under each condition.

### Fitting of the molecular structure of the Smc2/4 dimer to each AFM image

Next, we sought to extract the structure that best fits each AFM image from the structural ensembles generated by the coarse-grained molecular dynamics simulations. To do so, we first rotated the coarse-grained model so that the plane containing the representative amino acids of the heads and hinge was parallel to the AFM image. Each obtained configuration was flipped to obtain another configuration, which was treated as a separate one. Then, we projected the coordinates of each amino acid in the coarse-grained model onto the plane. In fitting other than the One-form images, the representative amino acid of the hinge was first superimposed on the geometric mean of the hinge pixels. Then, the structure was rotated around the hinge so that the Smc2 head overlapped with the head pixels. On the other hand, in fitting the One-form images, the geometric center of the two beads at the base of the arms was first superimposed on the geometric mean of the two arm pixels adjacent to the head pixels. Then, the structure was rotated so that the geometric center of the arm beads overlapped with the geometric mean of the arm pixels. After the fitting, we calculated the coverage score (the percentage of the arm pixels occupied by amino acids in the arm) as an index of how well the coarse-grained structure and the AFM image overlap. Please refer to Methods for details about detecting the head, hinge, and arm pixels from the AFM image and calculating the coverage score. Finally, we calculated the coverage score for each AFM image fitted using each structure from the simulation ensemble or the cryo-EM structure (PDB ID: 6YVU) (19) and selected the structure with the highest coverage score. Here, we analyzed 64 (Apo), 26 (AMP), 54 (ATPγS), 98 (ATP), and 74 (ADP) high quality images.

As a result, at least one structure with a coverage score of 0.7 or higher (more than 70% of the AFM arm pixels are occupied by amino acids of the generated structure) was identified for each AFM image, regardless of the presence or absence of ATP, ATPγS, AMP-PNP, and ADP (Figure 3A). Interestingly, when fitting the AFM images in the presence of ATP, ATPγS, and AMP-PNP, the coverage scores for 47%, 43%, and 43% of the AFM images became 0 (suitable structures not found) if the simulation results under the EO condition were removed from the structural ensemble (Figure 3B). Also, the coverage scores for 18%, 20%, and 46% of the AFM images became 0 if the simulation results under the DO condition were removed. On the other hand, the most significant change observed was only 20% in the fitting of the ATPγS state if the simulation results under the EC or DC condition were removed. These results suggested that the open hinge structure is important for fitting the AFM images in the presence of ATP, ATPγS, and AMP-PNP. When fitting AFM images in the presence of ADP, we found that excluding the simulation results of the EO and DO condition resulted in a coverage score of 0 in 18% and 68% of the AFM images, respectively (Figure 3B). These results suggested that the V-form with the open hinge and the disengaged heads is important for fitting AFM images in the presence of ADP.

**Figure 3.**
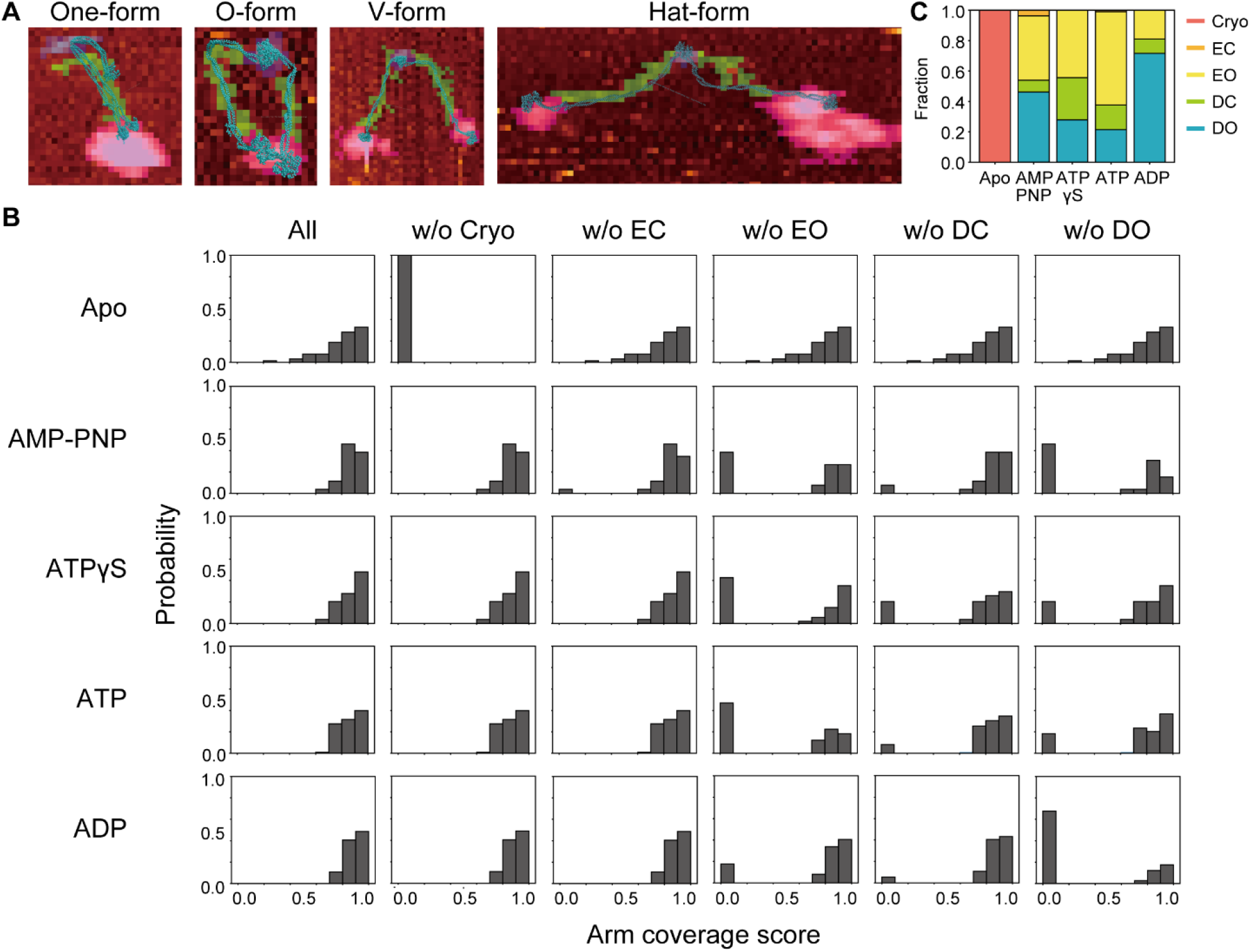
Fitting of the cryo-EM structure (PDB ID: 6YVU) (19) and the structures from the coarse-grained molecular dynamics simulations. (A) The representative fitted structures (light blue) on the target AFM images. The hinge, arm, and head pixels are colored purple, green, and pink, respectively. (B) The probability distributions of the arm coverage score defined in Methods (All). We also plotted the distribution when we removed the cryo-EM structure (w/o Cryo) (PDB ID: 6YVU) (19) or the structure from the coarse-grained simulations in the EC (w/o EC), EO (w/o EO), DC (w/o DC), and DO (w/o DO) conditions. (C) The fraction of the structural ensemble from which the fitted structure was selected. “Cryo” represents the probability of selecting the cryo-EM structure (PDB ID: 6YVU) (19) as the best fit.

When fitting the AFM images in the absence of nucleotides, we found the coverage score became 0 in 100% of the AFM images if the cryo-EM structure (19) was excluded (Fig. 3B). This result indicated that the AFM images in the absence of nucleotides resemble the One-form cryo-EM structure (19). In addition, the average elbow/head distance in the AFM images was almost identical to the cryo-EM structure as described above. Therefore, we concluded that we successfully observed the One-form structure, which was not observed in previous AFM studies, under the buffer conditions used in our solution AFM imaging.

Next, we plotted the statistics to clarify which structural ensemble best fits the AFM images in the presence and absence of ATP, ATPγS, AMP-PNP, and ADP (Figure 3C). In the absence of nucleotides, fitting with the One-form cryo-EM structure was always the best. In the presence of nucleotides, depending on individual AFM images, we found the best-fit structure derived from either the EO, DC, or DO ensemble. In particular, structures derived from the structural ensembles under EO and DO conditions accounted for most of the AFM images. This result suggested that it is necessary to assume an open hinge structure to fit the AFM images of the nucleotide-bound state. In the presence of ATP, ATPγS, and AMP-PNP, fitting with the structure derived from the structural ensemble under EO conditions was the best for an average of 49% of the AFM images. On the other hand, in the presence of ADP, fitting with the structure derived from the structural ensemble under EO conditions was the best for 19% of the AFM images. In addition, the proportion of DO+DC increased to 81% in the ADP state, suggesting that the ADP state induces a transition from an engaged to a disengaged state for the heads.

### Analysis of the fitted structural ensemble of the Smc2/4 dimer

To clarify how the condensin conformation in the presence and absence of ATP, ATPγS, AMP-PNP, and ADP were distributed in the structural space, we calculated eight features from each structure and performed principal component analysis (PCA) using the sklearn.decomposition library in Python. The calculated features are hinge angle, Smc2 elbow angle, Smc4 elbow angle, inter-elbow distance, inter-neck distance, inter-head distance, hinge-head distance, and inter-arm average distance (Refer to Methods for detailed definitions). PCA showed that 88% of the information was contained in up to the second principal component (PC2) (Figure 4A). Therefore, we reduced the dimensions to two for visualization and plotted them (Figure 4C; this plot includes structures in all solution conditions). As a result, we found that all structures were roughly divided into three clusters. The first cluster was for the single and isolated structure in the upper left corner of the PC plane (magenta), which is the One-form cryo-EM structure (19). The second cluster was for the structures in the lower left corner, which contain the O-form and I-form structures. The third cluster was for the structures in the upper right, which contains the V-form and Hat-form structures. In the presence of ATP, AMP-PNP, and ATPγS, the structures in the lower left (O-form and I-form) were enriched. On the other hand, in the presence of ADP, the structures in the upper right (V-form and Hat-form) were enriched. However, it was also found that the structures could not be classified entirely by the type of bound nucleotides. For example, in Figure 4C, the blue (ATP) and yellow dots overlap considerably. There are two possible explanations for this result, in addition to the possibility that the role of nucleotides is just to shift the population of the structures. The first possibility is nucleotide contamination. It has been known that commercially available ADP and AMP-PNP are contaminated with 0.1–2.0% ATP (35). The second possibility is the structural disruption due to adsorption to the AFM mica surface, accelerating the structural transition from the O-form to the V-form or Hat-form. The statistics of the number of conformations in each nucleotide state are explained below.

**Figure 4.**
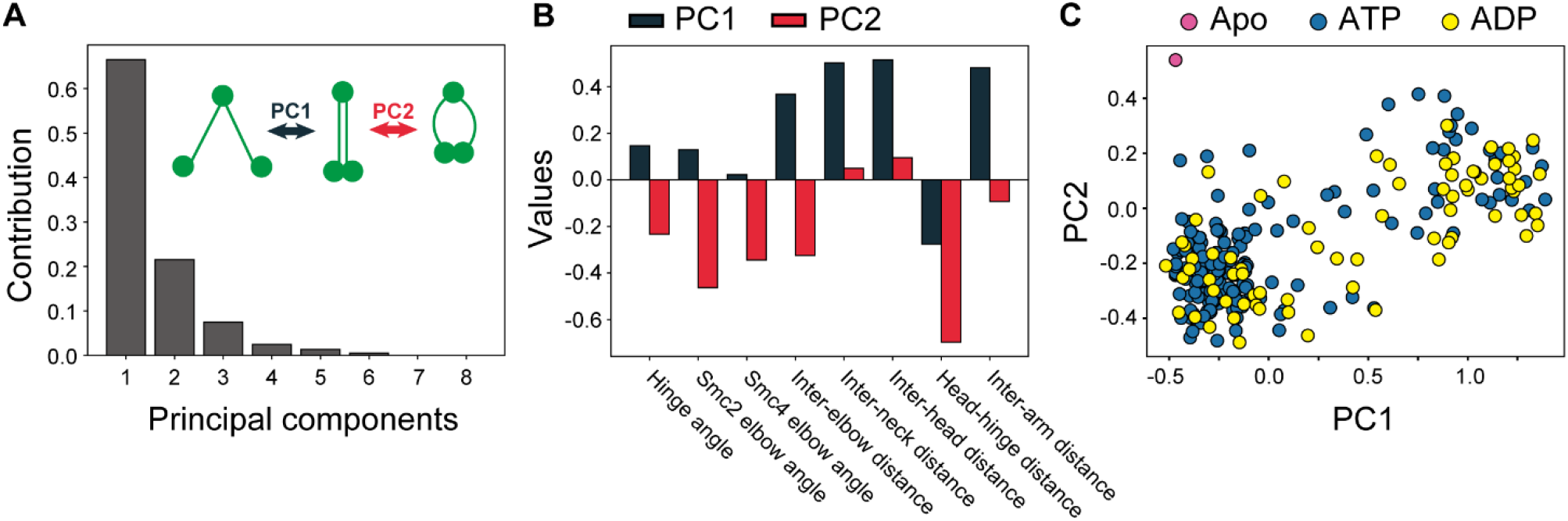
Principal component analysis of the structures fitted to each AFM image. (A) Contributions of each principal component. The inset shows the motion PC1 and PC2 roughly captured. (B) Contributions of each feature to PC1 (Dark blue) and PC2 (Red), respectively. (C) The features projected on the PC1-PC2 plane. “ATP” contains all the structures from the ATP, ATPγS, and AMP-PNP states.

It was found that the inter-elbow, hinge-head, inter-neck, and inter-arm average distances had significant contributions to PC1 and were positively correlated (Figure 4B). Therefore, PC1 is considered as an axis representing the transition dynamics between the I-form and V-form (Figure 4A inset). On the other hand, the inter-head distance was negatively correlated with the hinge angle, Smc2 elbow angle, Smc4 elbow angle, inter-elbow distance, and inter-arm average distance in PC2. Therefore, PC2 is considered as an axis representing the transition dynamics between the I-form and O-form (Figure 4A inset). Notably, in the presence of nucleotides, the structure was not enriched in the area where PC1 is small and PC2 is large (i.e., the upper left area of the PC plane in Fig. 4C). This result suggested that the decreased inter-head distance is linked to increased inter-arm distance and hinge angle and, therefore, that ATP-binding-induced head engagement is coupled to conformational changes in the hinge. It is consistent with a study by Gruber et al. on *Bacillus subtilis* Smc-ScpAB, in which ATP binding to the heads was shown to open the hinge via the arms using electron paramagnetic resonance (EPR) and cysteine cross-linking (36). In addition, we also found that after the elbow was unbent and the hinge was opened, the arms of yeast condensin were quite flexible and did not prevent the formation of either the I- or B-form.

Next, agglomerative clustering of the above eight feature vectors was performed to classify the structures further using Python’s sklearn.cluster library. First, the structures were broadly divided into Hat-form and others (Figure 5A). Then, they were divided into the One-form and others. Then, the structures were classified into V-form, I-form, and O-form. Interestingly, Hat-form and V-form were found to be classified into two separate clusters. And it was found that they also formed separate clusters on the PC plane shown in Fig. 4C (Fig. 5B). Furthermore, from the distance distribution between the two head domains (Figure 5C), we can see that the V-form formed a peak at a distance of 53Å, while the Hat-form formed a peak at a distance of 778Å, and there were few structures with distance in between. This result suggested that these conformations represent structural states with a significant free energy barrier for the transition between them. Excluding the structured domains at the N-terminus and C-terminus, the disordered linker of Brn1 contains approximately 500 amino acid residues. If the behavior of the disordered linker follows the random-flight polymer model, the average end-to-end distance is 85Å, significantly smaller than the 778 Å observed in the Hat-form. The probability of randomly extending to 778Å is diminishingly small (1 × 10^−44^). Considering these, together with previous studies (22), it is indicated that the Hat-form is formed by dissociation of the N-terminus of Brn1 from the Smc2 head due to ATP hydrolysis. In summary, these analyses revealed that condensin forms One-form, O-form, I-form, V-form, and Hat-form and that the V-form and Hat-form are distinct structural states.

**Figure 5.**
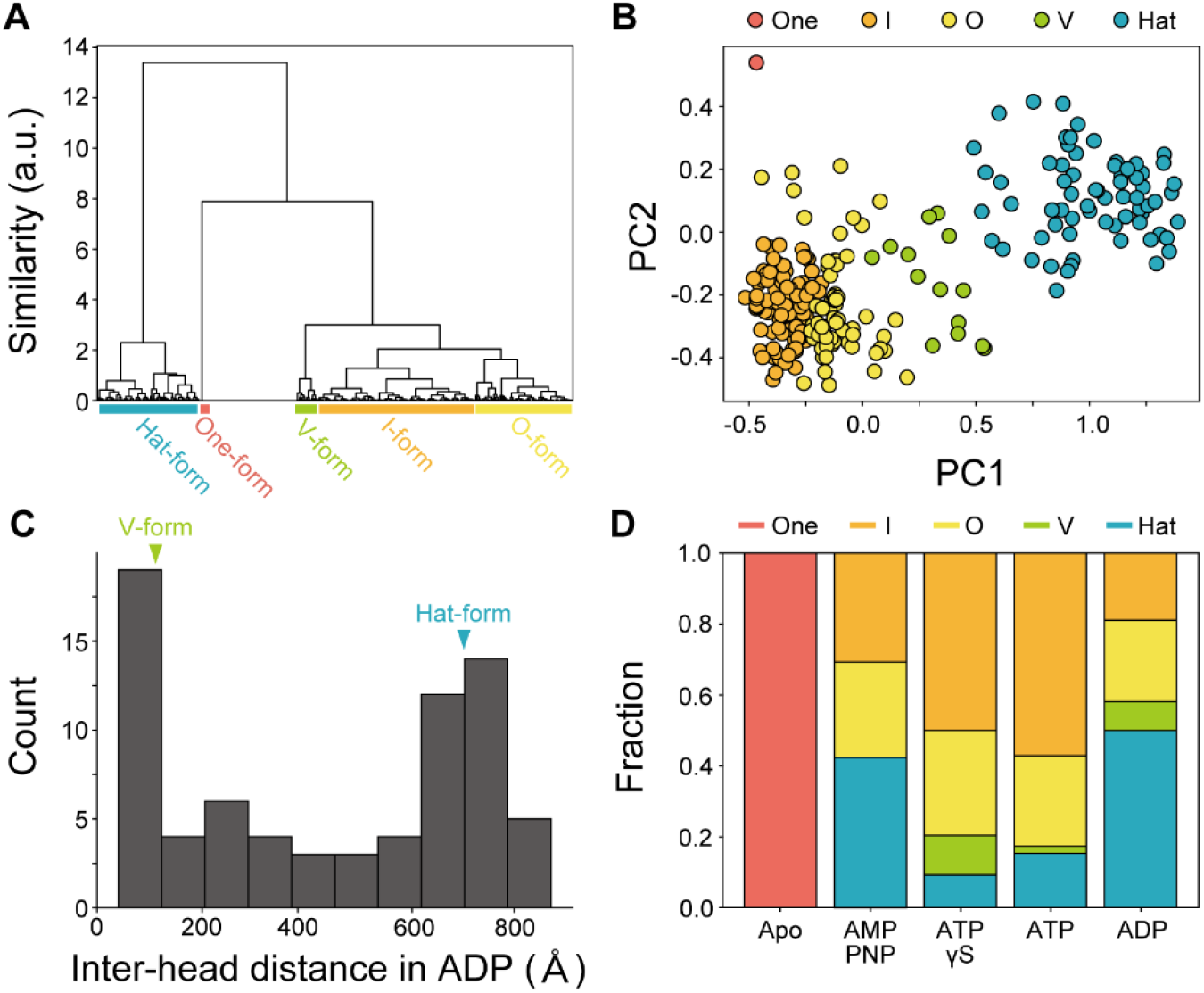
Analysis of the structural ensemble of the structures fitted to each AFM image. (A) The result of agglomerative clustering of the 8 features. (B) The same PC1-PC2 plane as Figure 4B in different coloring. (C) The histogram of the inter-head distance in the ADP state. The peak positions for the V- and Hat-forms are marked. (D) The fraction of the One-, I-, O-, V-, and Hat-forms in the presence and absence of AMP-PNP, ATPγS, ATP, and ADP.

Finally, we analyzed which conformations were enriched in each nucleotide solution condition (Figure 5D). First, the One-form population was 100% in the absence of nucleotides. In the presence of nucleotides, the O-form, I-form, V-form, and Hat-form were enriched. If we consider that the O-form and I-form are identical conformations that transition by arm fluctuation, the population of the O+I form was 73.3% on average in the ATP, AMP-PNP, and ATPγS states, while it was 41.9% in the ADP states. Instead, the V+Hat-form was 58.1% in the ADP state, which was higher than the average 26.7% in the ATP, AMP-PNP, and ATPγS states. These results supported that binding of ADP to the heads induces the transition from the engaged to the disengaged state. Contrary to expectations, the population of each conformation in the AMP-PNP solution was significantly different from that in either of the ATP/ATPγS solutions, rather resembling that in the ADP solution. The difference in molecular structure between AMP-PNP and ATP/ATPγS is thought to make AMP-PNP less effective at inducing head engagement. Interestingly, the Hat-form was enriched in the AMP-PNP and ADP states. This result is consistent with a previous study that ATP binding dissociates the N-terminus of Brn1 from the Smc2 head (22) and suggested that rebinding between Smc2 and Brn1 does not occur until the ADP release.

## DISCUSSION

This study used an integrated approach combining solution AFM imaging and coarse-grained molecular dynamics simulations to reveal nucleotide-dependent conformational changes in the yeast condensin complex. In the absence of nucleotides, the complex is known to adopt a One-form with a closed hinge, disengaged heads, and a bent elbow (19). This study revealed that the hinge transitions to the open state, and the heads transition to the engaged state, resulting in the O-form upon nucleotide binding. This result suggested that the transition to the engaged heads upon the ATP binding is linked to the transition to the open hinge structure. Furthermore, in the ADP-bound state, the Hat-form was mainly observed in which the hinge remains open, and the heads are disengaged. In the Hat-form, Smc2 and Brn1 are thought to be dissociated, and this result indicated that reassociation of the Brn1 and Smc2 heads occurs after ADP release.

Eeftens et al. performed high-speed AFM imaging to observe the dynamics of structural transitions between the V-, O-, B-, and P-forms of the Smc2/4 dimer in the absence of nucleotides (24). Ryu et al. also conducted high-speed AFM imaging to observe the dynamics of structural transitions between the O-, B-, and I-forms (possibly including the One-form) of the condensin holo-complex in the absence of nucleotides (25). Eeftens et al. could not observe the I-form, and Ryu et al. observed approximately 10% of the I-form. In this study, solution AFM imaging of the condensin holo-complex in the distinct buffer condition (previous study: 20 mM Tris-HCl pH 7.5, 50 mM NaCl, 2.5 mM MgCl_2_, 2.5 mM DTT; this study: 40 mM Tris-HCl pH 7.5, 125 mM KCl, 10 mM MgCl_2_, 1 mM DTT) revealed that the One-form was stable in the absence of nucleotides. This structure was similar to the cryo-EM structure without nucleotides (19). Considering the previous and current results, the One-form is thought to be stabilized by binding the accessory protein Ycs4 to the neck. We found an I-form, in which the elbow is no longer bent, as a conformation independent of the One-form. The transition from the One-form to the I-form is induced by nucleotide binding, and the dissociation of the hinge from the arm leads to flexible arm dynamics.

Vazquez Nunez et al. used EPR and cysteine cross-linking to show that the arms and hinges of prokaryotic SMCs open stepwise in the presence of nucleotides and double-stranded DNA (36). Meanwhile, the current study combined solution AFM imaging and coarse-grained molecular simulations to show that the arms and hinges of yeast condensin open in the presence of nucleotides. These two studies revealed that the coupling between the ATP-driven head engagement and hinge opening is conserved from prokaryotes to eukaryotes. In previous studies, the coupling could only be deduced from the distance distributions due to the method limitations (36). On the other hand, the current study used solution AFM imaging to directly visualize that the arms and hinges can open simultaneously, taking O-form. The result also revealed that the arm region flexibly opens and closes after ATP binding, transitioning between the O- and I-form. It will be necessary to comprehensively analyze the commonalities and specificities in the SMC complexes of prokaryotes and eukaryotes in the future.

Previously, the DNA segment capture model has been proposed as one of the molecular mechanisms of loop extrusion (37–39). In the first step of this model, ATP binding causes the arms and hinges to open, and the open hinge and the engaged heads bind to the base of the DNA loop. As mentioned above, the study by Vazquez Nunez et al. (36) and the current study revealed that ATP binding to the heads is linked to the structural transition to the open hinge. Our previous study predicted that the open hinge can bind to DNA using the coarse-grained molecular dynamics simulations (28). Therefore, our studies collectively suggested that the first step of the DNA segment capture model may be realized. Interestingly, amino acid substitutions on the surface exposed in the open hinge suppress the function of fission yeast condensin in cells, but the suppression is rescued by introducing an additional mutation into the kleisin subunit (40).

By atomic-level modeling and biochemical assays of the ATPase cycle of the condensin holo-complex, Hassler et al. showed that binding of one ATP induces dimerization of the SMC heads, whereas binding of two ATPs induces dissociation of the Brn1 N-terminus from the Smc2 head (22). In the current study, the Hat-form in which the hinge remains open and the heads are disengaged was predominantly observed in the presence of ADP. This result suggested that the Brn1 N-terminus remains dissociated from the Smc2 head even in the ADP-bound state. However, the ring compartments are dissolved when the Brn1 N-terminus dissociates from the Smc2 head while the heads are disengaged. Previous studies have shown that disruption of compartments is not essential for loop extrusion for human cohesin (41, 42) (not verified for budding yeast condensin). However, our result indicated that the molecular mechanism of loop extrusion must function even when the compartment is dissolved at a stage of the ATP hydrolysis cycle.

There are several limitations to this study. First, solution AFM imaging inevitably affects the molecular dynamics of the condensin molecules. In the presence of ATP, spontaneous structural transitions among the One-, O-, I-, V-, and Hat-form should occur, but the strong adsorption to the mica surface made it impossible to observe these transitions. Although structural changes were observed in the solution conditions of the previous study, the One-form could not be observed (25). Therefore, the surface and buffer conditions must be further examined. In addition, to reduce the computational cost and structural space in the coarse-grained molecular dynamics simulation, the Smc2/4 dimer was used to generate the structural ensemble for fitting AFM images, so the dynamics of the condensin holo-complex may not be accurately captured. In the future, simulations of the condensin holo-complex may enable us to capture the dynamics of the entire complex, including the accessory subunits. Despite these limitations, this study has combined experimental and computational methods to build a powerful framework for studying large-scale biomolecular assemblies and deepen our understanding of the molecular mechanisms of condensin.

## METHODS

### Solution AFM imaging

We purified condensin from budding yeast and performed solution AFM imaging the same way in our previous study (28). A cantilever for high-speed AFM imaging was attached to a tapping-mode high-speed atomic force microscope (spring constant ∼0.1 N/m, resonant frequency in water ∼0.5 MHz, quality factor in water ∼1.5). An imaging solution (40 mM Tris-HCl pH7.5, 125 mM KCl, 10 mM MgCl_2_, 1 mM DTT) containing 1 nM condensin and 0 mM or 4 mM nucleotides (Either of ATP, ADP, AMP-PNP, and ATPγS) was added to the mica stage. After 5 minutes of incubation, we washed the stage with the imaging solution and started the imaging. The free oscillation and set-point amplitude were set to 1–2 nm and 0.9 times that, respectively. An area of 150 nm × 150 nm was scanned.

### Detection of condensin in AFM images

In this study, we developed an algorithm to detect condensin molecules in AFM images. In this algorithm, we first correct the tilt and height of the mica surface. To accomplish this, we divide the AFM image into 5 pixels × 5 pixels windows. Then, we subtract the minimum height value from all pixels in the window. If these 25 pixels do not contain any pixels with a height value exceeding the minimum signal cutoff, we determine that the pixel in the center represents the mica surface. We then apply the least squares method to the surface pixels to correct the tilt and height of the surface. After the correction, pixels with height values exceeding the minimum signal cutoff are determined to be condensin pixels.

Next, the condensin pixels are classified into arm, hinge, and head pixels. To accomplish this, we calculate the volume (the sum of the height values) of a 5 pixels × 5 pixels area around the condensin pixels. If the volume is less than the hinge minimum volume cutoff, the condensin pixel is determined to be an arm pixel. Otherwise, it is determined to be a hinge/head pixel. Next, the remaining condensin pixels adjacent to arm and hinge/head pixels are classified into arm and hinge/head pixels, respectively, and this operation is repeated recursively to classify all the condensin pixels into arm and hinge/head pixel groups. Then, small pixel groups less than the pixel group size cutoff are discarded. The hinge/head pixel groups with a size less than the head minimum volume cutoff are classified as hinge pixel groups, and those with a size greater than the cutoff are classified as head pixel groups. The initial cutoffs were 9Å (minimum signal cutoff), 25 pixels (pixel group size cutoff), 120 nm^3^ (minimum hinge volume cutoff), and 738 nm^3^ (minimum head volume cutoff), but were adjusted ad hoc according to visual inspection.

Then, it is determined whether the arm, hinge, and head pixel groups constitute one molecule of condensin. First, a hinge pixel group is selected. Second, arm pixel groups within 2 pixels of the hinge pixel group are selected. If there are zero or three or more arm groups, the process starts with selecting the new hinge pixel group again. Third, it is checked whether head pixel groups exist within 2 pixels of the end of the arm pixel group opposite the hinge pixel group. If no such arm pixel groups exist, the process starts with selecting the new hinge pixel group. The selected hinge, arm, and head pixel groups are considered to constitute one molecule of condensin.

### Coarse-grained molecular dynamics simulations of the Smc2/4 dimer

We performed coarse-grained molecular dynamics simulations by integrating the Langevin equation with 0.3 CafemMol time unit (∼14.7 fs). The temperature and friction coefficient were set to 300K and 0.843, respectively. All simulations were performed using CafeMol 3.2 (33) (https://www.cafemol.org). A 4 × 10^8^ production run took 8.2 days on an Intel® Xeon® Gold 6326 CPU with a single core. Molecular structures were visualized using VMD 1.9.3 (43) (https://www.ks.uiuc.edu).

### Fitting coarse-grained structures to AFM images

We fitted the coarse-grained structures obtained by molecular dynamics simulations and the One-form cryo-EM structure (19) to the AFM image of condensin. First, the coarse-grained structures were rotated so that the plane spanned by three points (Smc2 I625 in the hinge, Smc2 V3 in the head, and Smc4 V232 in the head) was parallel to the AFM image. Here, we turned over the molecule and treated it as a separate molecule. Then, we projected the coarse-grained beads onto the plane. Next, in fitting other than the One-form cryo-EM structure (19), the geometric center of Smc2 A548 and Smc4 I648 was superimposed on the geometric mean of the hinge pixels. Then, the structure was rotated so the Smc2 head in the coarse-grained structure was on the head pixel group. If the Smc4 head was not on the head pixel group or the arms were not on the arm pixel group, the coverage score was set to 0. In fitting the One-form cryo-EM structure, the geometric center of Smc2 E204 and Smc4 L350 was superimposed on the geometric mean of the two arm pixels adjacent to the head pixel group. Then, the structure was rotated so that the geometric center of the arm beads (28 beads every 10th from Smc2 E194 and 28 beads every 10th from Smc4 S370) was superimposed on the geometric mean of the arm pixels.

Next, we calculated how many arm pixels were covered by the coarse-grained arm beads (28 beads every 10th from Smc2 E194 and 28 beads every 10th from Smc4 S370). We considered that the arm pixel was covered when the arm beads and the points dividing the segment between two beads into four resided in or next to the pixel. We then calculated the coverage score by dividing the number of covered arm pixels by the total number of arm pixels.

### Analysis of the structural ensemble of the Smc2/4 dimer

Eight features were calculated from each structure to analyze the structural ensemble of the Smc2/4 dimer obtained by the fitting. The first feature is the “hinge angle.” This is the angle formed by Smc2 L665, the geometric center of Smc2 A548 and Smc4 I648, and Smc4 E876, divided by *π*. The second is the “Smc2 elbow angle.” This is the angle formed by Smc2 K769, S787, and Q810, divided by *π*. The third is the “Smc4 elbow angle.” This is the angle formed by Smc4 Q564, I969, and L988, divided by *π*. The fourth is the “inter-elbow distance.” This is the distance between Smc2 S787 and Smc4 I969 divided by twice the distance between the hinge and elbow (Smc2 A548 and Smc2 S787; 14.67 nm) in the cryo-EM structure (PDB ID: 6YVU) (19). The fifth is the “Inter-neck distance.” This is the distance between Smc2 T225 and Smc4 K386 divided by twice the sum of the distances between the hinge and elbow and between the elbow and neck (Smc2 S787 and Smc2 T225; 23.2 nm) in the cryo-EM structure (PDB ID: 6YVU) (19). The sixth is the “Inter-head distance.” This is the distance between Smc2 Q1102 and Smc4 L299 divided by twice the sum of the distances between the hinge and elbow, between the elbow and neck, and between the neck and head (the distance between Smc2 T225 and Smc2 Q1102) in the cryo-EM structure (PDB ID: 6YVU) (19). The seventh is the “hinge-head distance.” This is the distance between the geometric center of Smc2 A548 and Smc4 I648, and the geometric center of Smc2 Q1102 and Smc4 L299 divided by twice the sum of the distances between the hinge and elbow, between the elbow and neck, and between the neck and head in the cryo-EM structure (PDB ID: 6YVU) (19). The eighth is the “inter-arm average distance.” This is the average distance of 28 beads every 10th from Smc2 E194 and 28 beads every 10th from Smc4 S370, divided by the maximum distance across all structures.

## DATA AVAILABILITY

The data that support the findings of this study are available from the corresponding author upon reasonable request.

## ACKNOWLEDGEMENTS

We would like to thank the laboratory members of the theoretical biophysics laboratory at Kyoto University for their discussions and assistance throughout this work. We also would like to thank Prof. Christian Haering for the discussions and Dr. Mayu S. Terakawa for purifying proteins. This work was supported by the Grant-in-Aid for Japan Society for the Promotion of Science Fellows (23KJ1265 to H.K.), by the Japan Science and Technology Agency (JST) grant (JPMJCR1762 to S.T.), by the Grant-in-Aid for Transformative Research Areas (24H00883; to T.T.), the grant from the Kyoto University Foundation (to T.T.), the grant from the Takeda Science Foundation (to T.T.), the grant from the Shimazu Science Foundation (to T.T.), the grant from the Inamori Foundation (to T.T.).

## AUTHOR CONTRIBUTIONS

H.K., S.T, and T.T. designed the project. H.K. performed the atomic force microscopy experiments with the assistance of N.K. and their analyses. H.K. also performed the coarse-grained molecular dynamics simulations and their analyses. H.K., N.K., S. T and T.T. co-wrote the manuscript.

## FIGURES

**Figure S1.**
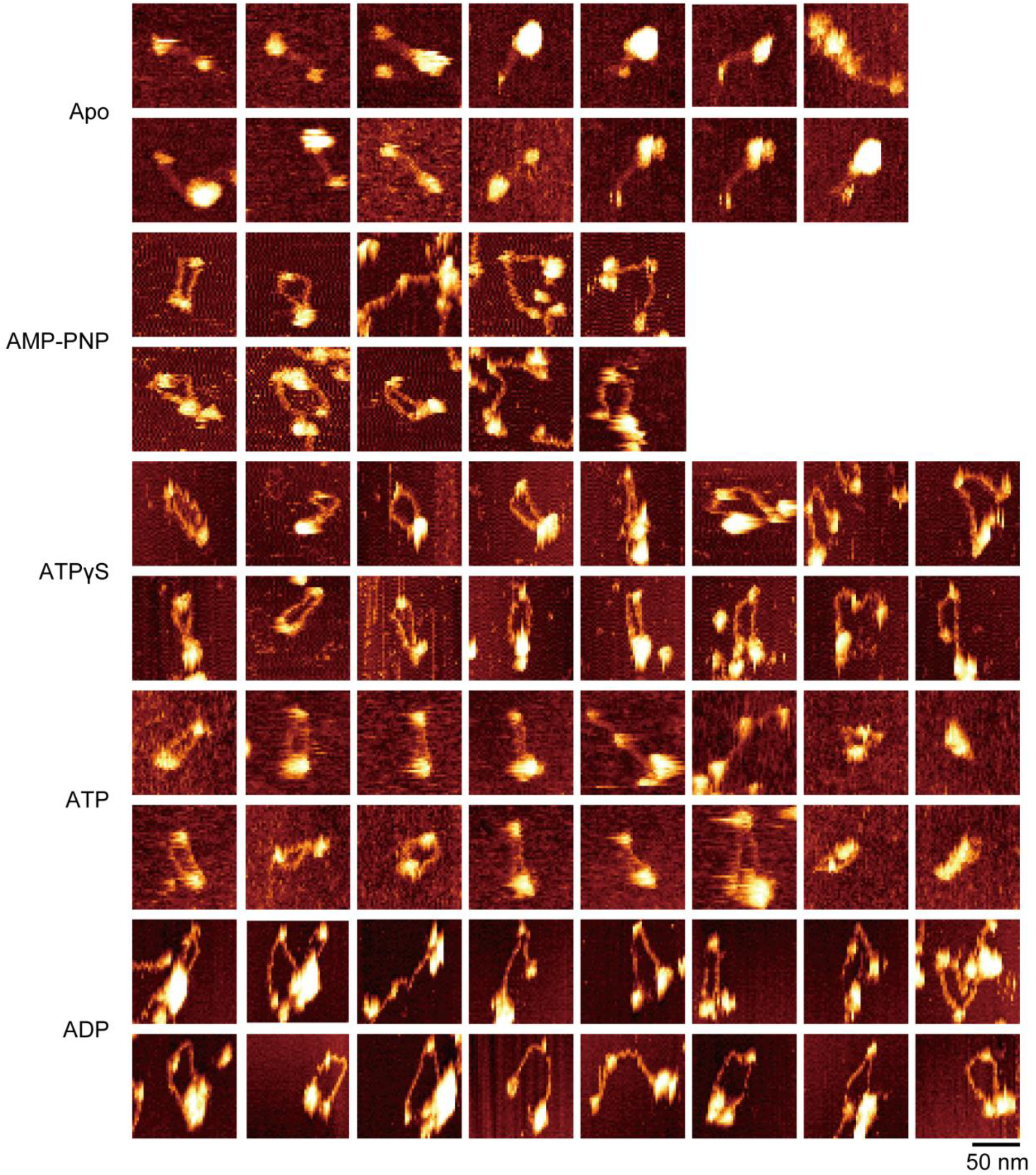
Example images of solution AFM imaging of budding yeast condensin holo-complex in the Apo (in the absence of nucleotides), AMP-PNP, ATPγS, ATP, and ADP states.

## REFERENCE

1. J. R. McIntosh, Mitosis. Cold Spring Harb Perspect Biol 8, a023218 (2016).

2. J. R. Paulson, D. F. Hudson, F. Cisneros-Soberanis, W. C. Earnshaw, Mitotic chromosomes. Seminars in Cell & Developmental Biology 117, 7–29 (2021).

3. N. Naumova, et al., Organization of the Mitotic Chromosome. Science 342, 948–953 (2013).

4. J. H. Gibcus, et al., A pathway for mitotic chromosome formation. Science 359, eaao6135 (2018).

5. K. Shintomi, T. S. Takahashi, T. Hirano, Reconstitution of mitotic chromatids with a minimum set of purified factors. Nat Cell Biol 17, 1014–1023 (2015).

6. N. L. Curtis, G. F. Ruda, P. Brennan, V. M. Bolanos-Garcia, Deregulation of Chromosome Segregation and Cancer. Annu. Rev. Cancer Biol. 4, 257–278 (2020).

7. T. Hirano, Condensin-Based Chromosome Organization from Bacteria to Vertebrates. Cell 164, 847– 857 (2016).

8. S. Yatskevich, J. Rhodes, K. Nasmyth, Organization of Chromosomal DNA by SMC Complexes. Annu. Rev. Genet. 53, 445–482 (2019).

9. I. F. Davidson, J.-M. Peters, Genome folding through loop extrusion by SMC complexes. Nat Rev Mol Cell Biol 22, 445–464 (2021).

10. A. Goloborodko, M. V. Imakaev, J. F. Marko, L. Mirny, Compaction and segregation of sister chromatids via active loop extrusion. eLife 5, e14864 (2016).

11. T. Terakawa, et al., The condensin complex is a mechanochemical motor that translocates along DNA. Science 358, 672–676 (2017).

12. M. Ganji, et al., Real-time imaging of DNA loop extrusion by condensin. Science 360, 102–105 (2018).

13. M. Kong, et al., Human Condensin I and II Drive Extensive ATP-Dependent Compaction of Nucleosome-Bound DNA. Molecular Cell 79, 99-114.e9 (2020).

14. J.-K. Ryu, et al., Condensin extrudes DNA loops in steps up to hundreds of base pairs that are generated by ATP binding events. Nucleic Acids Research 50, 820–832 (2022).

15. A. Pastic, et al., Chromosome compaction is triggered by an autonomous DNA-binding module within condensin. Cell Reports 43, 114419 (2024).

16. J. Park, J.-J. Kim, J.-K. Ryu, Mechanism of phase condensation for chromosome architecture and function. Exp Mol Med 56, 809–819 (2024).

17. T. Hirano, K. Kinoshita, SMC-mediated chromosome organization: Does loop extrusion explain it all? Current Opinion in Cell Biology 92, 102447 (2025).

18. K. Nasmyth, C. H. Haering, The structure and function of SMC and kleisin complexes. Annu. Rev. Biochem. 74, 595–648 (2005).

19. B.-G. Lee, et al., Cryo-EM structures of holo condensin reveal a subunit flip-flop mechanism. Nat Struct Mol Biol 27, 743–751 (2020).

20. X. Chu, J. Wang, Deciphering the molecular mechanism of the cancer formation by chromosome structural dynamics. PLoS Comput Biol 17, e1009596 (2021).

21. F. Bürmann, J. Löwe, Structural biology of SMC complexes across the tree of life. Current Opinion in Structural Biology 80, 102598 (2023).

22. M. Hassler, et al., Structural Basis of an Asymmetric Condensin ATPase Cycle. Molecular Cell 74, 1175-1188.e9 (2019).

23. B.-G. Lee, J. Rhodes, J. Löwe, Clamping of DNA shuts the condensin neck gate. Proc. Natl. Acad. Sci. U.S.A. 119, e2120006119 (2022).

24. J. M. Eeftens, et al., Condensin Smc2-Smc4 Dimers Are Flexible and Dynamic. Cell Reports 14, 1813–1818 (2016).

25. J.-K. Ryu, et al., The condensin holocomplex cycles dynamically between open and collapsed states. Nat Struct Mol Biol 27, 1134–1141 (2020).

26. Y.-M. Soh, et al., Molecular Basis for SMC Rod Formation and Its Dissolution upon DNA Binding. Molecular Cell 57, 290–303 (2015).

27. J. J. Griese, G. Witte, K.-P. Hopfner, Structure and DNA binding activity of the mouse condensin hinge domain highlight common and diverse features of SMC proteins. Nucleic Acids Research 38, 3454–3465 (2010).

28. H. Koide, N. Kodera, S. Bisht, S. Takada, T. Terakawa, Modeling of DNA binding to the condensin hinge domain using molecular dynamics simulations guided by atomic force microscopy. PLoS Comput. Biol. 17, e1009265 (2021).

29. D. J. Müller, A. Engel, The height of biomolecules measured with the atomic force microscope depends on electrostatic interactions. Biophysical Journal 73, 1633–1644 (1997).

30. W. Li, T. Terakawa, W. Wang, S. Takada, Energy landscape and multiroute folding of topologically complex proteins adenylate kinase and 2ouf-knot. Proc. Natl. Acad. Sci. USA 109, 17789–17794 (2012).

31. S. Takada, et al., Modeling Structural Dynamics of Biomolecular Complexes by Coarse-Grained Molecular Simulations. Acc. Chem. Res. 48, 3026–3035 (2015).

32. T. Terakawa, J. Higo, S. Takada, Multi-scale Ensemble Modeling of Modular Proteins with Intrinsically Disordered Linker Regions: Application to p53. Biophysical Journal 107, 721–729 (2014).

33. H. Kenzaki, et al., CafeMol: A Coarse-Grained Biomolecular Simulator for Simulating Proteins at Work. J. Chem. Theory Comput. 7, 1979–1989 (2011).

34. W. Kabsch, C. Sander, Dictionary of protein secondary structure: Pattern recognition of hydrogen-bonded and geometrical features. Biopolymers 22, 2577–2637 (1983).

35. N. Eswar, et al., Comparative Protein Structure Modeling Using Modeller. CP in Bioinformatics 15 (2006).

36. R. Vazquez Nunez, Y. Polyhach, Y.-M. Soh, G. Jeschke, S. Gruber, Gradual opening of Smc arms in prokaryotic condensin. Cell Reports 35, 109051 (2021).

37. J. F. Marko, P. De Los Rios, A. Barducci, S. Gruber, DNA-segment-capture model for loop extrusion by structural maintenance of chromosome (SMC) protein complexes. Nucleic Acids Research 47, 6956–6972 (2019).

38. S. K. Nomidis, E. Carlon, S. Gruber, J. F. Marko, DNA tension-modulated translocation and loop extrusion by SMC complexes revealed by molecular dynamics simulations. Nucleic Acids Research 50, 4974–4987 (2022).

39. M. Yamauchi, G. B. Brandani, T. Terakawa, S. Takada, SMC complex unidirectionally translocates DNA by coupling segment capture with an asymmetric kleisin path. [Preprint] (2024). Available at: http://biorxiv.org/lookup/doi/10.1101/2024.04.29.591782.

40. X. Xu, M. Yanagida, Suppressor screening reveals common kleisin–hinge interaction in condensin and cohesin, but different modes of regulation. Proc Natl Acad Sci USA 116, 10889–10898 (2019).

41. I. F. Davidson, et al., DNA loop extrusion by human cohesin. Science 366, 1338–1345 (2019).

42. B. Pradhan, et al., SMC complexes can traverse physical roadblocks bigger than their ring size. Cell Reports 41, 111491 (2022).

43. W. Humphrey, A. Dalke, K. Schulten, VMD: Visual molecular dynamics. Journal of Molecular Graphics 14, 33–38 (1996).

